# CFIm25-Dependent Alternative Polyadenylation in AKT2 mRNA Programs Macrophage Polarization

**DOI:** 10.1101/2025.06.19.660629

**Authors:** Srimoyee Mukherjee, Atish Barua, Marzieh Naseri, Claire L. Moore

## Abstract

Macrophage polarization is essential for immune responses, tissue homeostasis, and progression of many diseases. It is a tightly regulated process involving an intricate network of signaling pathways and control mechanisms at the level of transcription, alternative mRNA splicing, translation and mRNA stability. However, regulation through alternative mRNA polyadenylation (APA), remains poorly understood. This study explores the function of CFIm25, a key APA regulator, in macrophage polarization. Our findings show that CFIm25 overexpression drives M1 polarization, as evident from increased nitric oxide synthase activity, CD80 expression, and pro-inflammatory cytokine secretion, but dampens the M2 phenotype. Conversely, CFIm25 knockdown suppresses M1 traits and promotes M2 characteristics. Functionally, CFIm25 enhances phagocytosis, migration, and cancer cell inhibition. Mechanistically, CFIm25 favors proximal polyadenylation site usage of *AKT2* mRNA, increasing Akt2 protein levels to support M1 polarization. Blocking this site with an antisense oligonucleotide reduces Akt2 expression and M1 traits. These findings establish CFIm25 as a crucial regulator of macrophage identity, offering insights into RNA-based immune regulation and potential therapeutic targets.

**In brief:** CFIm25 drives macrophage polarization toward the M1 phenotype through specific regulatory pathways. The presence of CFIm25 profoundly shifts surface marker expression, enhancing M1 markers while suppressing M2 signatures. This extends to biochemical properties, where CFIm25 boosts nitric oxide and pro-inflammatory cytokine production while reducing anti-inflammatory mediators. Functionally, CFIm25 enhances phagocytosis, inflammatory migration, and cancer cell killing. Mechanistically, this orchestration of the M1 polarization program involves CFIm25’s regulation of *AKT2* mRNA alternative polyadenylation which increases Akt2 protein expression and amplifies NF-κB pathway activation, a central driver of M1 polarization.

**Highlights:**

- CFIm25 overexpression enhances M1 surface markers and biochemical properties including nitric oxide production and pro-inflammatory cytokine secretion.
- CFIm25 promotes M1 functional activities including phagocytosis, migration, and cancer-cytotoxic effects.
- Blocking the *AKT2* proximal polyadenylation site usage causes a decrease in Akt2 protein, suppressing M1 polarization phenotypes including NOS activity and cytokine profiles.
- The CFIm25-Akt2-NF-κB axis represents a novel target for macrophage polarization reprogramming.

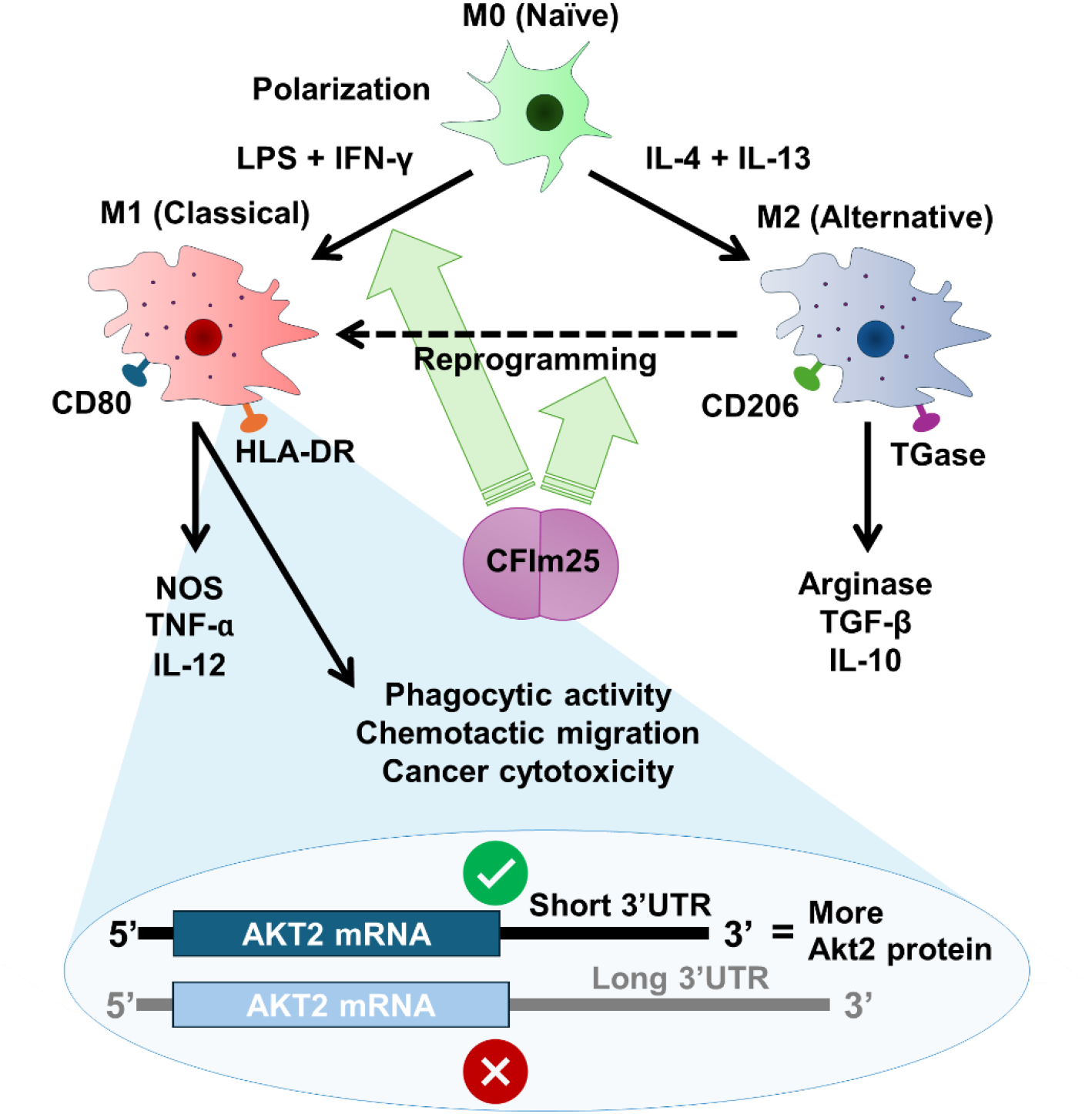

## Introduction

Macrophages are pivotal components of the innate immune system and exhibit a remarkable degree of plasticity in response to diverse environmental stimuli. This plasticity is most prominently exemplified by their capacity to polarize into distinct functional phenotypes, ranging along a spectrum from M1 (classically activated) to M2 (alternatively activated) macrophages ^1^. M1 macrophages are characterized by their pro-inflammatory properties, robust antimicrobial activities, and tumor-suppressive functions. These cells produce high levels of pro-inflammatory cytokines such as TNF-α, IL-1β, and IL-12, and exhibit enhanced antigen-presenting capabilities ^2^. In contrast, M2 macrophages display anti-inflammatory traits and are associated with wound healing, tissue repair, and, paradoxically, tumor promotion. M2 macrophages secrete anti-inflammatory mediators like IL-10 and TGF-β and play crucial roles in resolving inflammation and promoting tissue homeostasis ^3^. A major goal in immunotherapy is developing strategies to reprogram macrophage polarization states, as the ability to reliably shift macrophages between M1 and M2 phenotypes could enhance immune responses against various diseases. To attain this goal, it is necessary to understand molecular mechanisms that could serve as therapeutic targets for controlling macrophage fate and function.

The polarization of macrophages is a tightly regulated process involving an intricate network of signaling pathways and transcriptional control mechanisms. Key regulators of this process have been well characterized and include cytokines, kinases, transcription factors, and metabolic enzymes ^4^.

Mechanistic details of posttranscriptional controls at the level of alternative splicing, translation and mRNA stability are also being unraveled ^5^. However, the potential for regulation of macrophage polarization at the level of alternative polyadenylation (APA) remains largely unexplored. APA is a widespread mechanism of gene regulation that generates mRNA isoforms with different 3’ untranslated regions (UTRs) through the use of different polyadenylation sites. Changing the mRNA sequence in this way can profoundly impact mRNA stability, localization, translation efficiency, and protein coding capacity, ultimately influencing protein expression levels and cellular functions ^6–8^.

In recent years, CFIm25 (NUDT21, CPSF5), a key component of the Cleavage Factor Im (CFIm) complex, has emerged as a potent regulator of APA ^9^. Importantly, modulation of CFIm25 levels results in widespread changes in polyadenylation site selection, affecting the expression of numerous genes and, consequently, various cellular functions such as proliferation, cancer progression, and fibroblast differentiation ^10–13^. Of particular relevance to our current study, we have previously demonstrated that CFIm25 levels increase significantly during monocyte-to-macrophage differentiation, and its overexpression accelerates this process by affecting key factors including cell cycle regulators and the NF-κB signaling pathway ^14^. This finding established CFIm25 as a crucial regulator of macrophage development, where it influences the expression of differentiation-related genes through APA.

These observations set the stage for our current investigation into CFIm25’s impact on macrophage polarization. We hypothesized that CFIm25 might influence macrophage polarization through APA regulation. To test this hypothesis, we employed both gain- and loss-of-function approaches in THP-1 and HL-60 human monocytic cell lines, which are widely used models for studying macrophage biology ^15,16^. We examined the effects of CFIm25 modulation on M1 and M2 marker expression, cytokine production, utilization of arginine, and importantly, macrophage activities such as phagocytosis, migration and the ability to influence cancer cell growth, which are differentially regulated in M1 and M2 macrophages. In summary, CFIm25 promoted M1 phenotypes and suppressed those of M2-polarized cells. Our findings support a model in which CFIm25 can program macrophage fates through APA-mediated regulation of key genes, such as *AKT2*, which in turn modulates NF-κB signaling and its downstream inflammatory targets ^17,18^. This ability of CFIm25 to affect macrophage identity through APA represents a promising avenue for modulating immune responses in disease contexts where stronger antimicrobial or pro-inflammatory macrophage activity is desired, or alternatively, when inflammation needs to be suppressed.

## Results

### CFIm25 alteration drives surface marker expression in macrophages

To study macrophage polarization, we first established a reliable system to generate distinct M1 and M2 phenotypes. THP-1 and HL-60 monocytic cells were differentiated to naive macrophages (M0) and polarized to M1 subtypes with LPS and IFN-γ or to M2 phenotypes with IL-4 and IL-13. Flow cytometry analysis with dual staining for the M1 marker CD80 and M2 marker CD206 revealed clear differential expression patterns, with M1-polarized THP-1 cells showing predominant CD80 expression and M2-polarized cells exhibiting enhanced CD206 expression (Figure 1A-B). Similar patterns were observed in HL-60 cells (Figure S1A-B).

**Figure 1.**
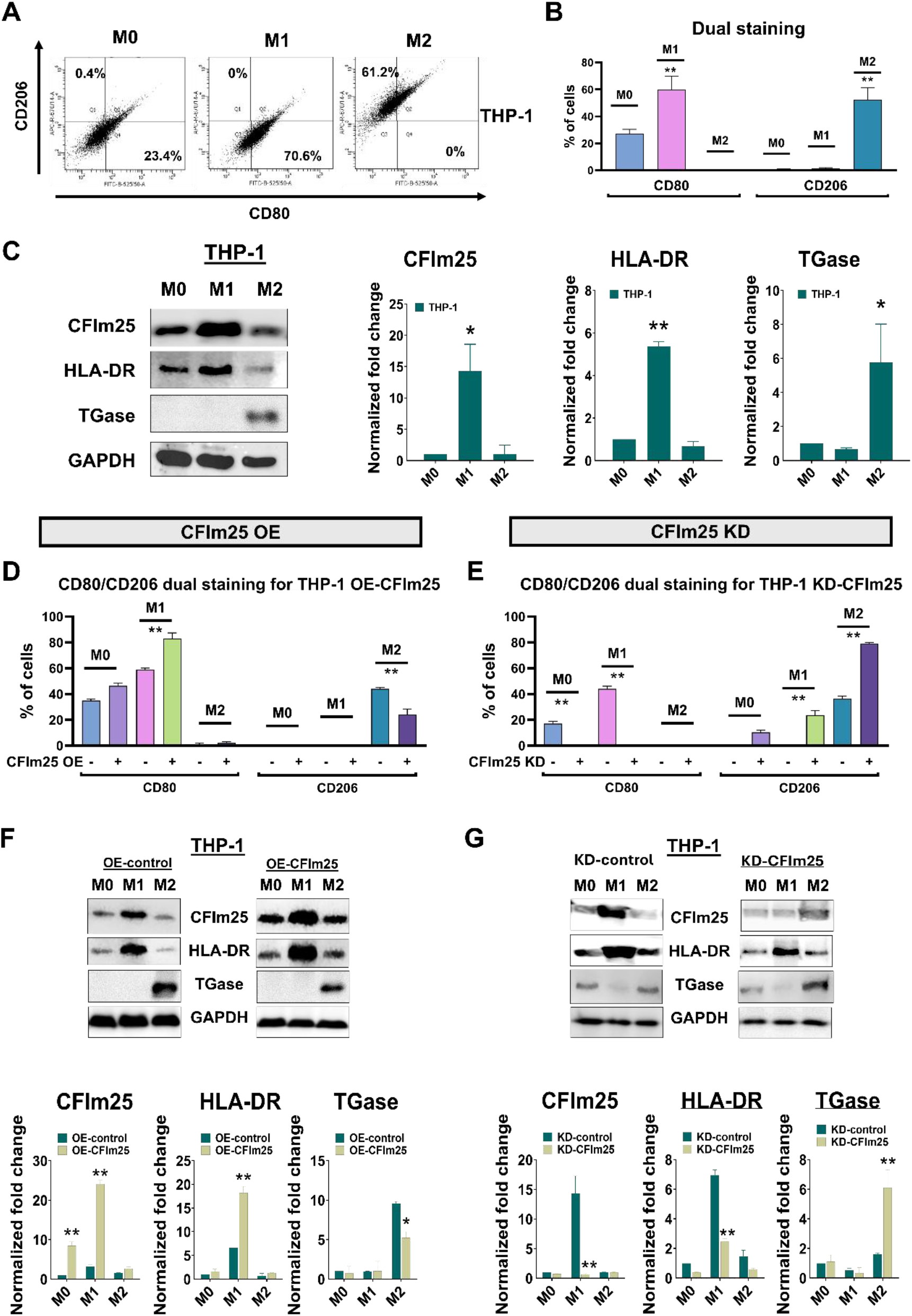
CFIm25 alteration drives macrophage surface marker expression in THP-1 cells. **(A)** Representative flow cytometry dot plots showing dual staining of M0, M1, and M2 polarized THP-1 cells for CD80 (M1 marker) and CD206 (M2 marker). Numbers indicate percentage of cells in each quadrant. **(B)** Quantification of flow cytometry data showing percentage of cells expressing CD80 and CD206 surface markers in M0, M1, and M2 polarized states. **(C)** Left: Western blot analysis of CFIm25, HLA-DR (M1 marker), and TGase (M2 marker) protein levels in M0, M1, and M2 polarized THP-1 cells. GAPDH serves as loading control. Right: Densitometric quantification of protein levels normalized to M0 state. **(D)** Flow cytometry analysis of CD80 and CD206 expression in CFIm25-overexpressing (OE) THP-1 cells under M0, M1, and M2 polarizing conditions. **(E)** Flow cytometry analysis of CD80 and CD206 expression in CFIm25-knockdown (KD) THP-1 cells under M0, M1, and M2 polarizing conditions. **(F)** Top: Western blot analysis comparing control and CFIm25-overexpressing THP-1 cells for expression of CFIm25, HLA-DR, and TGase under different polarization conditions. GAPDH serves as loading control. Bottom: Densitometric quantification of protein levels normalized to respective controls. **(G)** Top: Western blot analysis comparing control and CFIm25-knockdown THP-1 cells for expression of CFIm25, HLA-DR, and TGase under different polarization conditions. GAPDH serves as loading control. Bottom: Densitometric quantification of protein levels normalized to respective controls. For all experiments, data are normalized to respective controls and shown as mean ± SEM from three independent experiments. ***p < 0.001, **p < 0.01, *p < 0.05.

Western blot analysis for the M1 marker HLA-DR and M2 marker TGase provided additional validation of our polarization protocol in both THP-1 (Figure 1C) and HL-60 cells (Figure S1C). CFIm25 levels increased significantly in M1-polarized cells compared to naive macrophages but remained at M0 levels in M2-polarized cells. This dynamic regulation of CFIm25 expression during macrophage polarization suggested that CFIm25 might play a pivotal role in directing macrophages toward the M1 phenotype. This observation provided the fundamental rationale for our subsequent investigations into CFIm25’s function in macrophage polarization.

To directly test this hypothesis, we established stable cell lines with either CFIm25 overexpression or knockdown. Flow cytometry analysis showed that CFIm25 overexpression significantly enhanced CD80 expression in THP-1 M1 cells while reducing CD206 levels in M2 conditions (Figure 1D, raw data shown in S2A). Conversely, CFIm25 knockdown reduced CD80 expression in M0 and M1 cells while enhancing CD206 levels in both M1 and M2 cells (Figure 1E, raw data shown in S2B). Similar outcomes were observed in HL-60 cells (Figure S1D-E and raw data in S2C-D).

Western blot analysis corroborated these findings, with CFIm25 overexpression enhancing HLA-DR levels and reducing TGase expression in both THP-1 (Figure 1F) and HL-60 cells (Figure S1F), while CFIm25 knockdown produced the opposite effect (Figure 1G and S1G). Collectively, these results demonstrate that the differential expression of CFIm25 during macrophage polarization is not merely correlative but functionally significant, as experimental modulation of CFIm25 levels directly drives macrophage surface marker expression patterns across different cell lines.

### CFIm25 regulates macrophage biochemical properties

A distinguishing feature of M1 and M2 macrophages is their differential metabolism of arginine, with M1 cells producing nitric oxide via nitric oxide synthase (NOS) while M2 cells divert arginine toward arginase-dependent pathways. NOS and arginase assays revealed that M1-polarized macrophages displayed a significant increase in NOS activity accompanied by decreased arginase levels, while M2-polarized cells exhibited the opposite pattern in both THP-1 and HL-60 cells (Figure S3A-B, with representative flow cytometry plots shown in Figure S5A).

When we examined the impact of CFIm25 on these biochemical properties, we found that CFIm25 overexpression markedly enhanced NOS production in M1 cells and decreased arginase activity in M2 cells of both THP-1 (Figure 2A) and HL-60 lines (Figure S4A), with representative raw flow cytometry plots shown in Figure S5B. Conversely, CFIm25 knockdown led to decreased NOS activity in M1 cells and enhanced arginase activity in M2 cells (Figure 2B, Figure S4B, and representative raw flow cytometry plots in Figure S5C).

**Figure 2.**
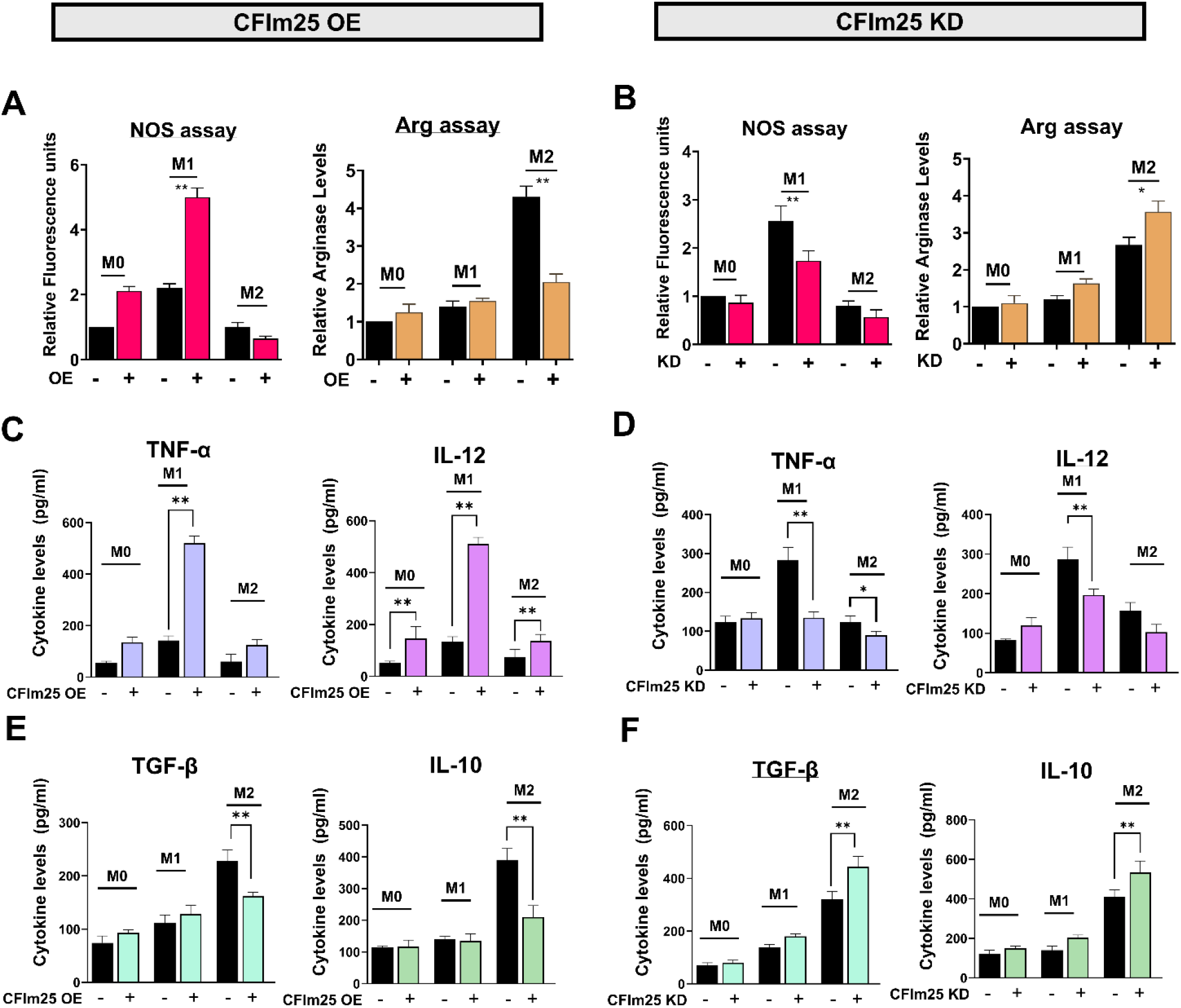
CFIm25 regulates macrophage biochemical properties in THP-1 cells. **(A)** Analysis of Nitric oxide synthase (NOS) and Arginase (Arg) activity in CFIm25-overexpressing (OE) THP-1 cells. Left: NOS activity measured by DAF-FM DA fluorescence in M0, M1, and M2 polarized cells with or without CFIm25 OE. Right: Arginase activity assay in the same conditions. **(B)** Analysis of NOS and arginase in CFIm25-knockdown (KD) HL-60 cells. Left: NOS activity measured by DAF-FM DA fluorescence in M0, M1, and M2 polarized cells with or without CFIm25 KD. Right: Arginase activity assay in the same conditions. For A-B, data are normalized to respective controls (M0 state) **(C)** ELISA measurement of M1 cytokines in CFIm25 OE cells. Left: TNF-α levels in M0, M1, and M2 conditions. Right: IL-12 levels in the same condition. **(D)** ELISA measurement of M1 cytokines in CFIm25 KD cells. Left: TNF-α levels in M0, M1, and M2 conditions. Right: IL-12 levels in the same condition. **(E)** ELISA measurement of M2 cytokines in CFIm25 OE cells. Left: TGF-β levels in M0, M1, and M2 conditions. Right: IL-10 levels in the same conditions. **(F)** ELISA measurement of M2 cytokines in CFIm25 KD cells. Left: TGF-β levels in M0, M1, and M2 conditions. Right: IL-10 levels in the same conditions. For C-F, data is represented as relative fluorescence units and are shown as mean ± SEM from three independent experiments **p < 0.01, *p < 0.05.

We next investigated cytokine production profiles, another key biochemical signature defining macrophage polarization states. As expected, M1-polarized control cells showed significantly higher levels of pro-inflammatory cytokines TNF-α and IL-12 compared to M2-polarized cells, while M2-polarized cells exhibited higher TGF-β and IL-10 levels in both cell lines (Figure S3C-F). CFIm25 overexpression significantly enhanced the production of M1 cytokines TNF-α and IL-12 in THP-1 cells (Figure 2C) and HL-60 cells (Figure S4C), while CFIm25 knockdown markedly reduced their levels (Figure 2D and Figure S4D).

Conversely, the M2-associated cytokines TGF-β and IL-10 were suppressed in CFIm25-overexpressing M2 cells (Figure 2E and Figure S4E) but elevated in CFIm25 knockdown cells (Figure 2F and Figure S4F). These results demonstrate that CFIm25 plays a pivotal role in regulating the biochemical properties that define macrophage polarization states.

### CFIm25 regulates core macrophage functions

Beyond markers, cytokines, and NOS activity, we investigated whether CFIm25 influences core macrophage functional activities. To evaluate phagocytic capacity, a hallmark of both M1 and M2 macrophages, we performed assays using fluorescently labeled IgG-coated beads, which mimic antibody-opsonized pathogens. Flow cytometry analysis revealed enhanced phagocytosis in CFIm25-overexpressing M0 and M1 THP-1 cells (Figure 3A), while upon CFIm25 knockdown, M1 cells displayed reduced phagocytic capacity (Figure 3B).

**Figure 3.**
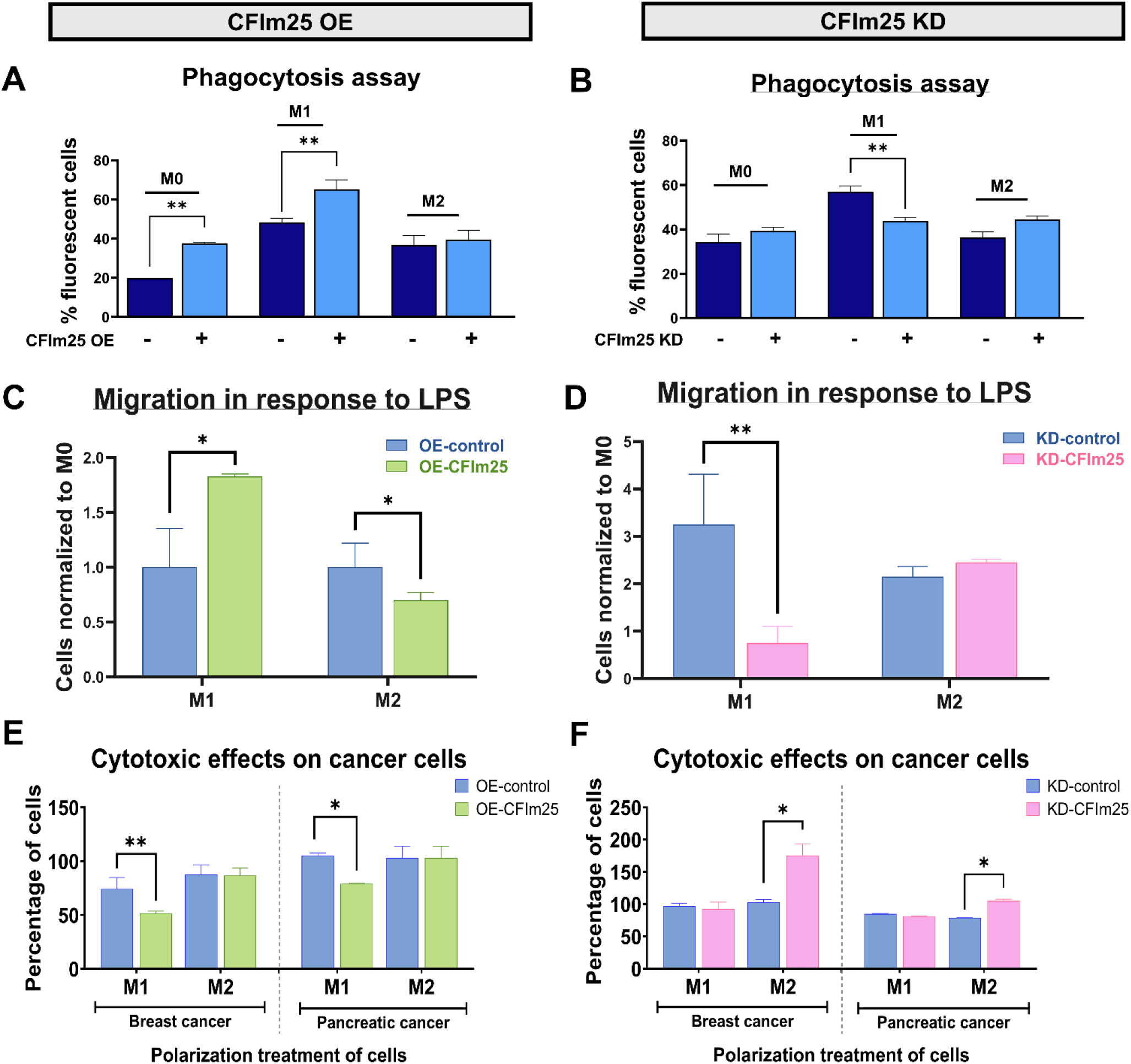
CFIm25 regulates core macrophage functions in THP-1 cells. Phagocytic activity in CFIm25-overexpressing (OE) THP-1 cells. Flow cytometry analysis of fluorescently labeled IgG-coated latex bead uptake in M0, M1, and M2 polarized cells with or without CFIm25 OE. Data shown as percentage of fluorescent cells, mean ± SEM from three independent experiments. **p < 0.01. **(B)** Phagocytic activity in M0, M1, and M2 polarized THP-1 cells with or without CFIm25 knockdown (KD). Data shown as percentage of fluorescent cells, mean ± SEM from three independent experiments. **p < 0.01. **(C)** Migration assay for CFIm25 OE cells. Quantification of cell migration through Transwell membranes in response to LPS. Data normalized to M0 control cells, mean ± SEM from three independent experiments. *p < 0.05. **(D)** Migration assay for CFIm25 KD cells. Data normalized to M0 control cells, mean ± SEM from three independent experiments. **p < 0.01, ***p < 0.001. **(E)** Cytotoxic activity of CFIm25-overexpressing macrophages against cancer cells. Indirect co-culture assay assessing the viability of breast cancer (MDA-MB-231) and pancreatic cancer (Mia-PaCa2) cells cultured in the lower chamber, with M1 or M2 polarized macrophages—with or without CFIm25 overexpression—seeded in the upper chamber separated by a permeable membrane. **(F)** Cytotoxic activity of CFIm25-overexpressing macrophages against cancer cells. Indirect co-culture assay assessing the viability of breast cancer (MDA-MB-231) and pancreatic cancer (Mia-PaCa2) cells cultured in the lower chamber, with M1 or M2 polarized macrophages—with or without CFIm25 knockdown—seeded in the upper chamber separated by a permeable membrane. Data are presented as percentage of viable cancer cells (mean ± SEM, n = 3 independent experiments). **p < 0.01, *p < 0.05.

Since migration is a hallmark property of M1 macrophages, which are activated and recruited in response to inflammatory cues such as lipopolysaccharide (LPS), a bacterial cell wall component that triggers Toll-like receptor 4 (TLR4) signaling and promotes a pro-inflammatory phenotype ^19^, we next evaluated the migratory capacity of CFIm25-modulated M1 THP-1 cells using a Boyden chamber assay. CFIm25-overexpressing M1 cells exhibited significantly enhanced migration toward LPS (Figure 3C), whereas knockdown cells demonstrated a reduced migratory response to the same stimulus (Figure 3D).

Finally, we investigated the effect of CFIm25 modulation on macrophages’ ability to influence cancer cell growth, another distinguishing feature between M1 and M2 phenotypes. In an indirect co-culture assay using a transwell system, CFIm25-overexpressing THP-1 M1 macrophages in the upper chamber inhibited growth of both breast cancer and pancreatic cancer cells compared to control (Figure 3E). Notably, CFIm25 knockdown did not alter the growth inhibitory effects of M1 macrophages but instead, resulted in an increase in cancer cell proliferation when M2 macrophages were present (Figure 3F). This suggests that CFIm25 not only enhances M1-mediated tumor suppression but also prevents M2 macrophages from fully exhibiting their pro-tumorigenic properties. These findings were mirrored in HL-60 cells as well (Figure S6), indicating a common mechanism in both macrophage models. These functional assays collectively demonstrate that CFIm25 orchestrates multiple core functional characteristics that define macrophage polarization states.

### CFIm25 affects polarization by alternative polyadenylation of *AKT2*

To elucidate the molecular mechanism underlying CFIm25’s effects on macrophage polarization, we investigated potential downstream targets regulated by APA. Our analysis focused on Akt2; a kinase previously established as promoting the M1 polarization state^20^. Analysis of *AKT2*’s gene structure revealed two distinct polyadenylation sites in its 3’ UTR, as illustrated in Figure 4A. Intriguingly, M1 macrophages predominantly utilized the proximal polyadenylation site, while M2 macrophages preferentially employed the distal site (Figure 4B). This differential usage of polyadenylation sites had functional consequences: M1 macrophages exhibited increased *AKT2* mRNA expression and corresponding elevated protein levels, while M2 macrophages showed reduced expression at both mRNA and protein levels (Figure 4B).

**Figure 4.**
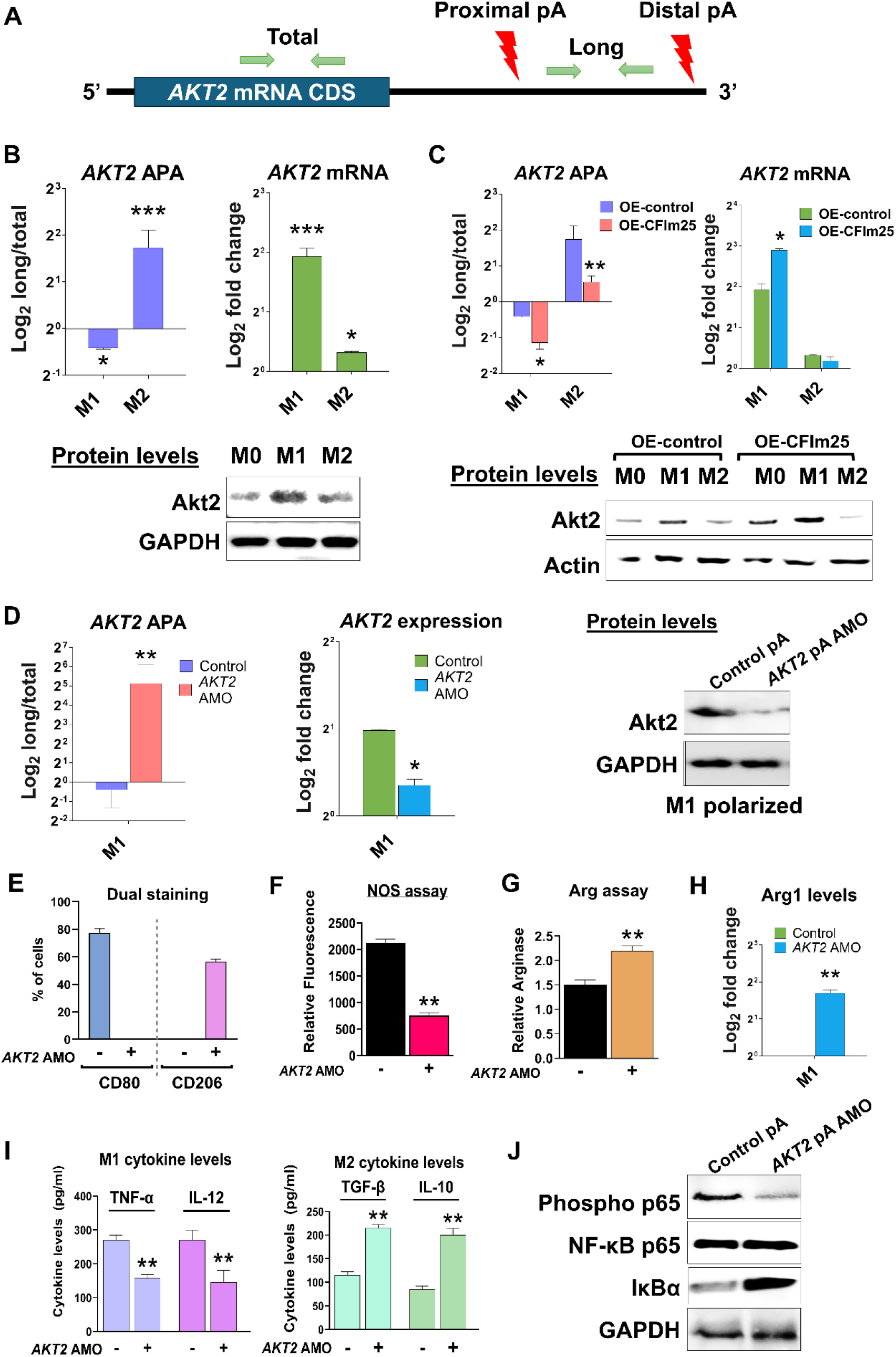
CFIm25 affects polarization by alternative polyadenylation of *AKT2*. **(A)** Schematic representation of *AKT2* mRNA showing the coding sequence (CDS), positions of proximal and distal polyadenylation (pA) sites (red lightning bolts), and locations of primer pairs used for total and long isoform detection (green arrows). **(B)** Analysis of AKT2 in M1 and M2 polarized THP-1 cells. Left: APA analysis showing Log2 ratio of long/total AKT2 mRNA. Right: Log2 fold change in total AKT2 mRNA levels. Bottom: Western blot showing Akt2 protein levels, with GAPDH as loading control. **(C)** Impact of CFIm25 overexpression on Akt2. Left: APA analysis showing Log2 ratio of long/total AKT2 mRNA in control and CFIm25 OE cells. Right: Log2 fold change in total AKT2 mRNA levels. Bottom: Western blot showing Akt2 protein levels, with Actin as loading control. **(D)** Effect of AKT2 proximal polyadenylation site blockade using antisense morpholino oligonucleotides (AMO). Left: APA analysis showing Log2 ratio of long/total AKT2 mRNA. Middle: Log2 fold change in total AKT2 mRNA levels. Right: Western blot showing Akt2 protein levels in M1 polarized cells, with GAPDH as loading control. Data for A-D are normalized to respective controls **(E)** Effect of AKT2 pA AMO on macrophage surface markers by dual staining. Figure shows quantification of flow cytometry data showing percentage of cells expressing CD80 and CD206 surface markers in M1 polarized cells with or without AKT2 pA AMO **(F)** NOS activity measured by DAF-FM DA fluorescence in M1 polarized cells with or without AKT2 AMO. **(G)** Arginase activity assay in M1 polarized cells with or without AKT2 AMO. **(H)** Arginase gene expression by qRT-PCR in M1 polarized cells with or without AKT2 AMO. **(I)** Impact of AKT2 AMO on cytokine production. Left: M1 cytokines (TNF-α and IL-12) levels. Right: M2 cytokines (TGF-β and IL-10) levels measured by ELISA For all experiments, data is shown as mean ± SEM from three independent experiments. ***p < 0.001, **p < 0.01, *p < 0.05. **(J)** Western blot analysis of NF-κB pathway components in M1 polarized cells treated with control or AKT2 pA AMO for phospho-p65, total NF-κB p65 and IκB-α where GAPDH serves as loading control.

CFIm25 overexpression further accentuated this pattern, promoting enhanced usage of the proximal polyadenylation site and resulting in increased *AKT2* mRNA and protein levels in M1 cells (Figure 4C). To determine the functional significance of this APA-mediated regulation, we designed an antisense Morpholino oligomer (AMO) specifically targeting the *AKT2* proximal polyadenylation site. Treatment with this AMO effectively blocked proximal site usage, forcing utilization of the distal site, which resulted in transcript lengthening and decreased *AKT2* mRNA and protein levels (Figure 4D).

The functional consequences of *AKT2* APA manipulation were profound and phenocopied the effects observed with CFIm25 knockdown. AMO-mediated lengthening of *AKT2* transcripts led to a striking decrease in cells expressing the M1 marker CD80, with an increase in expression of the M2 marker CD206 as observed by dual staining (Figure 4E, raw data shown in S7A). The AMO treatment also decreased NOS activity (Figure 4F, raw data shown in S7B) and elevated arginase levels (Figure 4G). In line with this, qPCR analysis showed a significant upregulation of *ARG1* expression following *AKT2* AMO treatment (Figure 4H), consistent with a previous report that Akt2 activity suppresses activation of the Arg1 promoter ^21^. Moreover, cytokine profiling revealed that *AKT2* AMO treatment reduced M1-associated cytokines such as TNF-α and IL-12, while increasing M2 cytokines like TGF-β and IL-10 (Figure 4I).

Akt2 has been implicated as an activator of the NF-kB signaling pathway^17^. Consistent with this role, AMO-induced lengthening of *AKT2* mRNAs and the resulting decrease in Akt2 protein led to a marked reduction in phosphorylated NF-*κ*B p65 (Ser536), accompanied by increased levels of its cytoplasmic inhibitor, IκB-α (Figure 4J). IκB-α functions by sequestering NF-κB dimers in the cytoplasm, thereby preventing their nuclear translocation and transcriptional activation of pro-inflammatory target genes, including TNF-α, IL-12, and iNOS^4^. These results establish a mechanistic pathway whereby CFIm25-mediated APA regulation of Akt2 promotes M1 polarization through an enhanced NF-κB signaling axis, unveiling a novel post-transcriptional mechanism governing macrophage inflammatory responses.

## Discussion

In this report, we identify CFIm25 as a critical regulator of macrophage polarization. Our findings demonstrate that CFIm25 overexpression promotes M1 polarization by enhancing M1 phenotypic markers, increasing pro-inflammatory cytokine and NOS production, and boosting key M1 functions such as phagocytosis, migration, and cancer cell killing while suppressing M2 characteristics. Conversely, CFIm25 knockdown shifts macrophages towards the M2 phenotype. Mechanistically, CFIm25 regulates Akt2 expression through APA, promoting proximal polyadenylation site usage that increases Akt2 protein levels, thereby enhancing NF-κB signaling and M1 polarization.

Reprogramming macrophage polarization represents a promising strategy for immunotherapy in conditions ranging from cancer to chronic inflammation and autoimmune disorders ^22–24^. Enhanced M1 polarization can boost anti-tumor immunity and pathogen clearance, while M2 polarization promotes tissue repair and resolution of inflammation ^1,2,4^. However, taking advantage of macrophage plasticity will require precise molecular tools to control macrophage phenotypes in disease contexts. Current strategies largely focus on transcriptional and signaling mechanisms, including modulation of cytokine receptors, manipulation of STAT-family transcription factors such as STAT1 and STAT6 (which function as both signal transducers and transcriptional regulators in cytokine signaling pathways), metabolic reprogramming, and epigenetic modification ^2^.^4^. Our findings introduce post-transcriptional regulation through APA as a novel and potentially effective mechanism for macrophage reprogramming.

APA affects most mammalian genes ^25^ and can significantly influence protein expression through changes in protein coding sequence, mRNA stability, localization, and translation efficiency ^6-8,26^. However, the contribution of APA to macrophage polarization has remained elusive. Wilton et al. demonstrated that pro-inflammatory polarization of primary human macrophages leads to widespread changes in APA, affecting genes important for immune function ^27^. While they identified APA alterations, they did not establish functional consequences of this APA. Consistent with our findings and supporting the conservation of a CFIm25-mediated regulatory mechanism, Chen et al. recently reported in late 2024 that CFIm25 (Nudt21) was upregulated in mouse bone marrow-derived macrophages (BMDMs) after stimulation with the M1 inducers IFNγ and LPS ^28^. They also found that myeloid-specific CFIm25 depletion in a mouse model reduced M1 cytokine expression by disrupting APA of autophagy regulators and conferred a protective effect against colitis in mice, further underscoring the role of CFIm25-mediated APA in macrophage function.

Our study demonstrates that CFIm25 programming of macrophage polarization affects not only M1-specific cytokine production, but also multiple other functional hallmarks of both M1 and M2 phenotypes. This regulation can be achieved either through direct CFIm25 modulation or via targeted manipulation of *AKT2* APA. By promoting proximal polyadenylation site usage in *AKT2* transcripts, CFIm25 increases Akt2 protein levels, which in turn specifically supports M1 polarization ^21^. The NF-κB pathway serves as a central regulator of inflammatory responses in macrophages, controlling the expression of over 100 target genes involved in immunity ^29^. Akt2 enhances NF-κB signaling by activating the IKK complex, which phosphorylates and targets IκB-α for degradation, thereby releasing NF-κB dimers for nuclear translocation. Concurrently, Akt2 phosphorylates the p65 (RelA) subunit, promoting transcriptional activation of key inflammatory genes ^17,18^. Our data reveal that CFIm25-mediated modulation of *AKT2* APA, a regulatory axis not previously reported, directly alters NF-κB signaling, as evidenced by changes in p65 phosphorylation, IκB-α accumulation, and the expression of canonical NF-κB target genes.

The CFIm25-Akt2-NF-κB axis represents a promising target for modulating macrophage-driven processes in various diseases. In cancer immunotherapy, enhancing CFIm25 expression in macrophages could promote M1 polarization to boost anti-tumor immunity. Conversely, in inflammatory disorders, targeting CFIm25 or *AKT2* APA could help resolve excessive inflammation by shifting macrophages toward an M2 phenotype. The precise nature of gene-specific APA regulation also offers the potential for interventions with fewer off-target effects compared to strategies targeting broad transcriptional regulators.

## Limitations of the study

Several important considerations should be acknowledged in our study. While we demonstrate CFIm25’s role in regulating macrophage plasticity, CFIm25 likely represents just one component of a broader polarization-specific APA regulatory network in macrophages, with other RNA-binding proteins potentially playing complementary roles. For example, knock-down of SRSF12, an RNA binding protein upregulated during M1 polarization, caused APA changes in several genes in primary human macrophages^27^, but functional consequences of the SRSF12 depletion were not reported. Future work is also needed to address how APA during macrophage polarization is coordinated with other RNA processing steps such as splicing and base modification. In addition, the therapeutic targeting of APA remains technically challenging because unlike the *AKT2* proximal polyadenylation site, the regions flanking many polyadenylation sites are A/U-rich and designing AMOs against a specific site is not always possible. Finally, macrophage-targeted delivery of CFIm25 mRNA, siRNAs against CFIm25, or *AKT2* AMOs, using nanocarriers such as those designed by Zhang, et al.^27^, may be needed for precise in vivo manipulation of macrophage polarization in immune-related diseases.

## Lead Contact and Materials Availability

Further information and requests for resources and reagents should be directed to and will be fulfilled by the Lead Contact, Claire L. Moore.

## Author contributions

CM and SM conceived the study, with CM providing overall project oversight. SM and AB developed strategy and methodology, designed the experiments, acquired the data, and organized the findings. MN contributed to the design of the Morpholino oligonucleotide by conducting the initial BLAST analyses, validating candidate sequences, and optimizing the AMO transfection conditions. All authors interpreted the results. The first draft of the manuscript was written by SM, and all authors contributed to manuscript revisions. SM handled correspondence and final submission. All authors approved the submitted version.

## Acknowledgements

This work was supported by the National Institutes of Health (NIH) grant 1R01AI152337 (to C.M.). The authors also acknowledge the Tufts Flow Cytometry Core for technical support.

## Experimental Model and Subject Details Cell culture and treatment

All cells were maintained in RPMI 1640 medium supplemented with 2 mM L-glutamine and 10% heat-inactivated fetal bovine serum (FBS) at 37°C in a humidified atmosphere of 5% CO2. Differentiation of the HL-60 (ATCC® CCL-240) and THP-1 (ATCC® TIB-202) human monocytic cell lines into macrophages was performed by treatment with 3 nM phorbol-12-myristate-13-acetate (PMA; Sigma-Aldrich) for up to 24 hours. Fresh media was added following that and after two days, polarization towards the M1 phenotype was induced by treating differentiated naïve macrophages with 50 ng/ml lipopolysaccharide (LPS) and 10 ng/ml Interferon-γ (IFN-γ) for a further 48 hours. Polarization towards the M2 phenotype was induced by culturing naive macrophages with 25 ng/ml each of interleukin-4 (IL-4) and interleukin-13 (IL-13) for 48 hours.

## Method Details

### NOS and arginase assays

Nitric oxide (NO) production was measured using the cell-permeable fluorescent indicator DAF-FM DA (MilliporeSigma™). THP-1 and HL-60 cells seeded in 96-well plates (1 × 10^5^ cells/well), were either M0 (PMA-differentiated) for 24h or polarized to M1 or M2 states for another 48h. Then they were washed with PBS and incubated with 5 μM DAF-FM DA for 30 minutes at 37°C in the dark. Following incubation, cells were washed twice with phosphate-buffered saline (PBS), and fluorescence intensity was measured using flow cytometry. Arginase activity was determined using the Arginase Activity Assay Kit (Sigma-Aldrich), which measures the conversion of arginine to urea and ornithine. The urea produced reacts with the kit substrate to generate a colored product proportional to arginase activity. Briefly, cells were lysed in 100 μL of lysis buffer containing protease inhibitors. Following centrifugation at 10,000g for 10 minutes, the supernatant was collected, and arginase activity was measured according to the manufacturer’s protocol. The colorimetric reaction was measured at 430 nm and normalized to total protein content.

### Dual staining and flow cytometry

For surface marker analysis, THP-1 and HL-60 cells (M0, M1, or M2 polarized) were collected and washed twice with cold PBS containing 10% FBS. Cells (1 × 10^6^) were stained with FITC or PE anti-human CD80 Antibody (BioLegend) and APC Mouse Anti-Human CD206 (BD Pharmingen™) overnight at 4°C in the dark. Following incubation, cells were washed twice with PBS/10% FBS to remove unbound antibodies. Flow cytometric analysis was performed on an Attune flow cytometer (Thermo Fisher Scientific), and data were analyzed using FlowJo software (Tree Star Inc.). Gates were set using appropriate isotype control antibodies, and at least 10,000 events were recorded for each sample.

### Western blot analysis

Western blot was performed with total cell extracts of naïve and polarized THP-1 and HL-60 cells. Briefly, 50-80 µg protein was separated on a 10% polyacrylamide-SDS or bis-tris gel and transferred to PVDF membrane. The membrane was blocked with 5% milk followed by rocking overnight at 4°C with primary antibodies (antibody details provided in Supplementary Table 1). On the next day, after three washes, the membrane was incubated with an HRP-conjugated secondary antibody for 1 hour, developed with SuperSignal™ West Pico Chemiluminescent Substrate (Thermo Fisher Scientific), and visualized with a gel doc. β-actin or GAPDH was used as the loading control and pre-stained protein markers were used as internal molecular mass standards. Each western blot was performed in three biological replicates. Western blot quantification was performed using ImageJ with normalization to loading controls and PMA-treated controls.

### Lentivirus construction and cell treatment

The CFIm25 OE lentivirus vector was a kind gift from Shervin Assassi (University of Texas Health Science Center at Houston, TX). It was generated by cloning the coding sequence (CDS) of human CFIm25 into the pLV-EF1a-IRES-Puro Vector (Addgene) that contains an EF-1a promoter upstream of an IRES element to co-express the puromycin marker. The CFIm25 CDS was inserted between the EF-1a and IRES ^30^. The vector with an insertion of GFP was used as a control. The CFIm25-overexpressing and control lentiviruses were then generated using the Dharmacon packaging mix and monocytic cells were transfected with CFIm25 OE or control virus. For CFIm25 knockdown, shRNA directed to its coding region was cloned in the LT3-GEPIR vector, viruses were made and transfected into monocytic cells to produce stable cell lines.

### ELISA for cytokines

The BioLegend® Legend Max™ kit was used to perform ELISA according to manufacturer’s protocol. It is a sandwich ELISA, in which specific monoclonal antibodies against the human M1 cytokines TNF-α and IL-12 and the M2 cytokines IL-10 and TGF-β are precoated on a 96-well strip-well plate. Supernatants from naïve and M1 or M2 polarized HL-60 and THP-1 cells are used to quantify the amount of expressed cytokines and represented as picogram/ml.

### Migration assay

THP-1 cells were seeded on a Transwell™ insert (8 µm pore; Corning, NY, USA) and incubated at 37°C, with 5% CO2. Following that, PMA only or M1 and M2 inducers were added as mentioned before and incubated for 24 hours. Then, these inserts were placed into a chamber containing RPMI medium with 100 ng/mL LPS to create a migratory stimulus for M1 macrophages. Transwell cultures were incubated for an additional 48 h to allow migration onto the other side of the semi-permeable membrane. Cells that migrated across the membrane were then counted with the help of Trypan blue staining and represented as bar graphs.

### Phagocytosis assay

The phagocytic activity was determined using a phagocytosis assay kit (Cayman Chemicals), in accordance with the manufacturer’s instructions. Briefly, the cells were seeded on a 96-well plate at a density of 2 × 10^5^ cells/well, differentiated into macrophages and polarized, with or without CFIm25 overexpression or knockdown. Subsequently, latex beads coated with fluorescently labeled IgG-FITC complex (Sourcee) were added to each well. After incubation for 2 hours, the cells were washed with assay buffer, and the fluorescence intensity was measured by flow cytometry.

### Indirect coculture assay for cytotoxic effects on cancer cells

THP-1 and HL-60 monocytes were seeded on the insert of a transwell system. Following that, PMA only or M1 and M2 inducers were added as mentioned before and incubated for 72 hours. MDA-MB-231 and Mia-Paca2 cells were seeded in the lower chamber of another transwell at a density of 1 × 10^5^ cells/ml. After the polarization was complete and the cancer cells grew, these inserts were placed into a chamber containing cancer cell lines. After 48 hours of polarization and coculture, the cancer cells in the lower chamber were assessed for viability using the MTT assay (Sigma) to evaluate the cytotoxic effects of secreted factors on cancer cells.

### RT-qPCR analysis for APA and expression

RNA isolation was performed for naïve and polarized THP-1 and HL-60 cells as described earlier. Briefly, total RNA extraction was carried out on 2×10^6^ cells using Trizol according to the manufacturer’s protocol. 1.5 µg of RNA was subjected to reverse transcription using oligo dT primer and Superscript III reverse transcriptase. The cDNA was amplified by qPCR with specific primers (Supplementary Table 2) using the C1000™ thermal cycler with CFX96 Touch Real-Time PCR Detection System (Bio-Rad Laboratories). The relative expression of genes was analyzed quantitatively by the ΔΔC method. ACTB RNA was used as the normalization control for RT-qPCR-based RNA expression analyses because its level did not change upon differentiation. Primers for total or long AKT2 transcripts were designed according to the p(A) site annotations featured in the poly(A)_DB database ^31^.

### Antisense oligonucleotide treatment

A custom Morpholino antisense oligonucleotide was designed to block the proximal polyadenylation (pA) site of the Akt2 mRNA, preventing recruitment of the polyadenylation machinery. Following Gene Tools, LLC guidelines for targeting RNA-binding sites, sequences flanking the proximal pA signal were extracted and analyzed to identify an optimal binding site. The Morpholino sequence, 5′-TGAGTGTCTTTATTGCTTGTACCGT-3′, was selected to hybridize directly over the pA signal (underlined) and its immediate surrounding region for use at 37°C. Specificity was verified via NCBI BLAST to minimize off-target hybridization within critical regions such as 5′ untranslated regions (5′ UTRs), splice junctions, or alternative PASs where binding could potentially alter gene expression, and secondary structure at the target site was assessed to ensure accessibility. The oligo was synthesized by Gene Tools at a 300 nmol scale, resuspended in sterile nuclease-free water upon arrival, and stored at –20°C. A standard control Morpholino with a sequence 5’-CCTCTTACCTCAGTTACAATTTATA-3’, that targets a human beta-globin intron mutation causing beta-thalassemia, was used in parallel to control for nonspecific effects.

For delivery, THP-1 cells were cultured in complete medium containing up to 10% serum. Differentiation was performed by treatment with PMA and the following day, AMOs in fresh media were added to a final concentration of 10 μM/mL, followed by the addition of Endo-Porter PEG (Gene Tools, LLC) at a final concentration of 6 μM/mL. Then polarization towards the M1 phenotype was induced for a further 48 hours, and experiments were performed in triplicate.

### Quantification and statistical analysis

All experiments were performed in at least three independent sets. Data are presented as mean ± standard error of the mean (SEM). Statistical analysis was performed using GraphPad Prism 6.01. Student’s t-test was used to determine the significance between groups. Statistical significance was indicated as: * = P ≤ 0.05; ** = P ≤ 0.01; *** = P ≤ 0.001; **** = P ≤ 0.0001.

## References

1. Italiani, P., and Boraschi, D. (2014). From Monocytes to M1/M2 Macrophages: Phenotypical vs. Functional Differentiation. Front Immunol 5, 514. 10.3389/fimmu.2014.00514.

2. Shapouri-Moghaddam, A., Mohammadian, S., Vazini, H., Taghadosi, M., Esmaeili, S.A., Mardani, F., Seifi, B., Mohammadi, A., Afshari, J.T., and Sahebkar, A. (2018). Macrophage plasticity, polarization, and function in health and disease. J Cell Physiol 233, 6425–6440. 10.1002/jcp.26429.

3. Wang, S., Wang, J., Chen, Z., Luo, J., Guo, W., Sun, L., and Lin, L. (2024). Targeting M2-like tumor-associated macrophages is a potential therapeutic approach to overcome antitumor drug resistance. NPJ Precis Oncol 8, 31. 10.1038/s41698-024-00522-z.

4. Wang, N., Liang, H., and Zen, K. (2014). Molecular mechanisms that influence the macrophage m1-m2 polarization balance. Front Immunol 5, 614. 10.3389/fimmu.2014.00614.

5. Guillemin, A., Kumar, A., Wencker, M., and Ricci, E.P. (2021). Shaping the Innate Immune Response Through Post-Transcriptional Regulation of Gene Expression Mediated by RNA-Binding Proteins. Front Immunol 12, 796012. 10.3389/fimmu.2021.796012.

6. Mitschka, S., and Mayr, C. (2022). Context-specific regulation and function of mRNA alternative polyadenylation. Nat Rev Mol Cell Biol 23, 779–796. 10.1038/s41580-022-00507-5.

7. Zhang, Q., and Tian, B. (2023). The emerging theme of 3’UTR mRNA isoform regulation in reprogramming of cell metabolism. Biochem Soc Trans 51, 1111–1119. 10.1042/BST20221128.

8. Gallicchio, L., Olivares, G.H., Berry, C.W., and Fuller, M.T. (2023). Regulation and function of alternative polyadenylation in development and differentiation. RNA Biol 20, 908–925. 10.1080/15476286.2023.2275109.

9. Hardy, J.G., and Norbury, C.J. (2016). Cleavage factor Im (CFIm) as a regulator of alternative polyadenylation. Biochem Soc Trans 44, 1051–1057. 10.1042/BST20160078.

10. Masamha, C.P. (2023). The emerging roles of CFIm25 (NUDT21/CPSF5) in human biology and disease. Wiley Interdiscip Rev RNA 14, e1757. 10.1002/wrna.1757.

11. Sun, X., Li, J., Sun, X., Liu, W., and Meng, X. (2020). CFIm25 in Solid Tumors: Current Research Progress. Technol Cancer Res Treat 19, 1533033820933969. 10.1177/1533033820933969.

12. Weng, T., Ko, J., Masamha, C.P., Xia, Z., Xiang, Y., Chen, N.Y., Molina, J.G., Collum, S., Mertens, T.C., Luo, F., et al. (2019). Cleavage factor 25 deregulation contributes to pulmonary fibrosis through alternative polyadenylation. J Clin Invest 129, 1984–1999. 10.1172/JCI122106.

13. Ghosh, S., Ataman, M., Bak, M., Borsch, A., Schmidt, A., Buczak, K., Martin, G., Dimitriades, B., Herrmann, C.J., Kanitz, A., and Zavolan, M. (2022). CFIm-mediated alternative polyadenylation remodels cellular signaling and miRNA biogenesis. Nucleic Acids Res 50, 3096–3114. 10.1093/nar/gkac114.

14. Mukherjee, S., Barua, A., Wang, L., Tian, B., and Moore, C.L. (2025). The alternative polyadenylation regulator CFIm25 promotes macrophage differentiation and activates the NF-kappaB pathway. Cell Commun Signal 23, 115. 10.1186/s12964-025-02114-1.

15. Gupta, D., Shah, H.P., Malu, K., Berliner, N., and Gaines, P. (2014). Differentiation and characterization of myeloid cells. Curr Protoc Immunol 104, 22F 25 21–22F 25 28. 10.1002/0471142735.im22f05s104.

16. Qin, Z. (2012). The use of THP-1 cells as a model for mimicking the function and regulation of monocytes and macrophages in the vasculature. Atherosclerosis 221, 2–11. 10.1016/j.atherosclerosis.2011.09.003.

17. Dan, H.C., Cooper, M.J., Cogswell, P.C., Duncan, J.A., Ting, J.P., and Baldwin, A.S. (2008). Akt-dependent regulation of NF-kappaB is controlled by mTOR and Raptor in association with IKK. Genes Dev 22, 1490–1500. 10.1101/gad.1662308.

18. Bai, D., Ueno, L., and Vogt, P.K. (2009). Akt-mediated regulation of NFkappaB and the essentialness of NFkappaB for the oncogenicity of PI3K and Akt. Int J Cancer 125, 2863–2870. 10.1002/ijc.24748.

19. Wu, T.T., Chen, T.L., and Chen, R.M. (2009). Lipopolysaccharide triggers macrophage activation of inflammatory cytokine expression, chemotaxis, phagocytosis, and oxidative ability via a toll-like receptor 4-dependent pathway: validated by RNA interference. Toxicol Lett 191, 195–202. 10.1016/j.toxlet.2009.08.025.

20. Vergadi, E., Ieronymaki, E., Lyroni, K., Vaporidi, K., and Tsatsanis, C. (2017). Akt Signaling Pathway in Macrophage Activation and M1/M2 Polarization. J Immunol 198, 1006–1014. 10.4049/jimmunol.1601515.

21. Arranz, A., Doxaki, C., Vergadi, E., Martinez de la Torre, Y., Vaporidi, K., Lagoudaki, E.D., Ieronymaki, E., Androulidaki, A., Venihaki, M., Margioris, A.N., et al. (2012). Akt1 and Akt2 protein kinases differentially contribute to macrophage polarization. Proc Natl Acad Sci U S A 109, 9517–9522. 10.1073/pnas.1119038109.

22. Gordon, S., and Taylor, P.R. (2005). Monocyte and macrophage heterogeneity. Nat Rev Immunol 5, 953–964. 10.1038/nri1733.

23. Mantovani, A., Sica, A., and Locati, M. (2005). Macrophage polarization comes of age. Immunity 23, 344–346. 10.1016/j.immuni.2005.10.001.

24. Murray, P.J. (2017). Macrophage Polarization. Annu Rev Physiol 79, 541–566. 10.1146/annurev-physiol-022516-034339.

25. Wang, R., Zheng, D., Yehia, G., and Tian, B. (2018). A compendium of conserved cleavage and polyadenylation events in mammalian genes. Genome Res 28, 1427–1441. 10.1101/gr.237826.118.

26. Di Giammartino, D.C., Nishida, K., and Manley, J.L. (2011). Mechanisms and consequences of alternative polyadenylation. Mol Cell 43, 853–866. 10.1016/j.molcel.2011.08.017.

27. Wilton, J., de Mendonca, F.L., Pereira-Castro, I., Tellier, M., Nojima, T., Costa, A.M., Freitas, J., Murphy, S., Oliveira, M.J., Proudfoot, N.J., and Moreira, A. (2023). Pro-inflammatory polarization and colorectal cancer modulate alternative and intronic polyadenylation in primary human macrophages. Front Immunol 14, 1182525. 10.3389/fimmu.2023.1182525.

28. Chen, Y., Chen, B., Li, J., Li, H., Wang, G., Cai, X., Zhang, Q., Liu, X., Kan, C., Wang, L., et al. (2024). Alternative mRNA polyadenylation regulates macrophage hyperactivation via the autophagy pathway. Cell Mol Immunol 21, 1522–1534. 10.1038/s41423-024-01237-8.

29. Liu, T., Zhang, L., Joo, D., and Sun, S.C. (2017). NF-kappaB signaling in inflammation. Signal Transduct Target Ther 2, 17023–. 10.1038/sigtrans.2017.23.

30. Weng, T., Huang, J., Wagner, E.J., Ko, J., Wu, M., Wareing, N.E., Xiang, Y., Chen, N.Y., Ji, P., Molina, J.G., et al. (2020). Downregulation of CFIm25 amplifies dermal fibrosis through alternative polyadenylation. J Exp Med 217. 10.1084/jem.20181384.

31. Wang, R., Nambiar, R., Zheng, D., and Tian, B. (2018). PolyA_DB 3 catalogs cleavage and polyadenylation sites identified by deep sequencing in multiple genomes. Nucleic Acids Res 46, D315–D319. 10.1093/nar/gkx1000.

