## Supplementary information for "CFIm25-Dependent Alternative Polyadenylation in AKT2 mRNA Programs Macrophage Polarization"

### Supplementary tables and figures:

### Supplementary Table 1. Antibodies used in this study:

*For western blot:*

| Catalogue no. | Name | Host | MW | Vendors |
|---|---|---|---|---|
| sc-81109 | CFIm25 | Mouse monoclonal | 26 kDa | Santa Cruz Biotechnology |
| sc-55593 | HLA-DR | Mouse monoclonal | 34 kDa | Santa Cruz Biotechnology |
| sc-48387 | TGase | Mouse monoclonal | 77 kDa | Santa Cruz Biotechnology |
| sc-365062 | GAPDH | Mouse monoclonal | 37 kDa | Santa Cruz Biotechnology |
| sc-81148 | Akt2 | Mouse monoclonal | 56 kDa | Santa Cruz Biotechnology |
| sc-47778 | β-actin | Mouse monoclonal | 43 kDa | Santa Cruz Biotechnology |
| 3033S | Phospho NF-κβ-p65 | Rabbit monoclonal | 65 kDa | Cell Signaling Technologies |
| 8242S | NF-κβ-p65 | Rabbit monoclonal | 65 kDa | Cell Signaling Technologies |
| sc-373893 | IκBα | Mouse monoclonal | 41 kDa. | Santa Cruz Biotechnology |
| 422301 | Human TruStain FcX™ | Monoclonal Antibody | $F_c$ blocker | Biolegend |

*For flow cytometry:*

| Catalogue no. | Name | Host | Fluorophore | Vendor |
|---|---|---|---|---|
| 305208 | CD80 | Mouse | PE | Biolegend |
| 321120 | CD206 | Mouse | APC | Biolegend |

### Supplementary Table 2. Primers used in this study:

| Primer name | Forward primer 5'-3' | Reverse primer 5'-3' |
|---|---|---|
| Akt2 total | CCAGTCCATCACAATCACACCC | GCCTGAAGAAGAACTGGAAAGGG |
| Akt2 long | TAGCCTGGATGTGTCTGGGC | GCTGGGGGAGGTGTTCCATC |
| Arg1 expression | TGATGTTGACGGACTGGACC | ATCTAATCCTGAGAGTAGCCCTGT |

**Supplementary Figure 1. CFIm25 alteration drives macrophage surface markers in HL-60 cells**

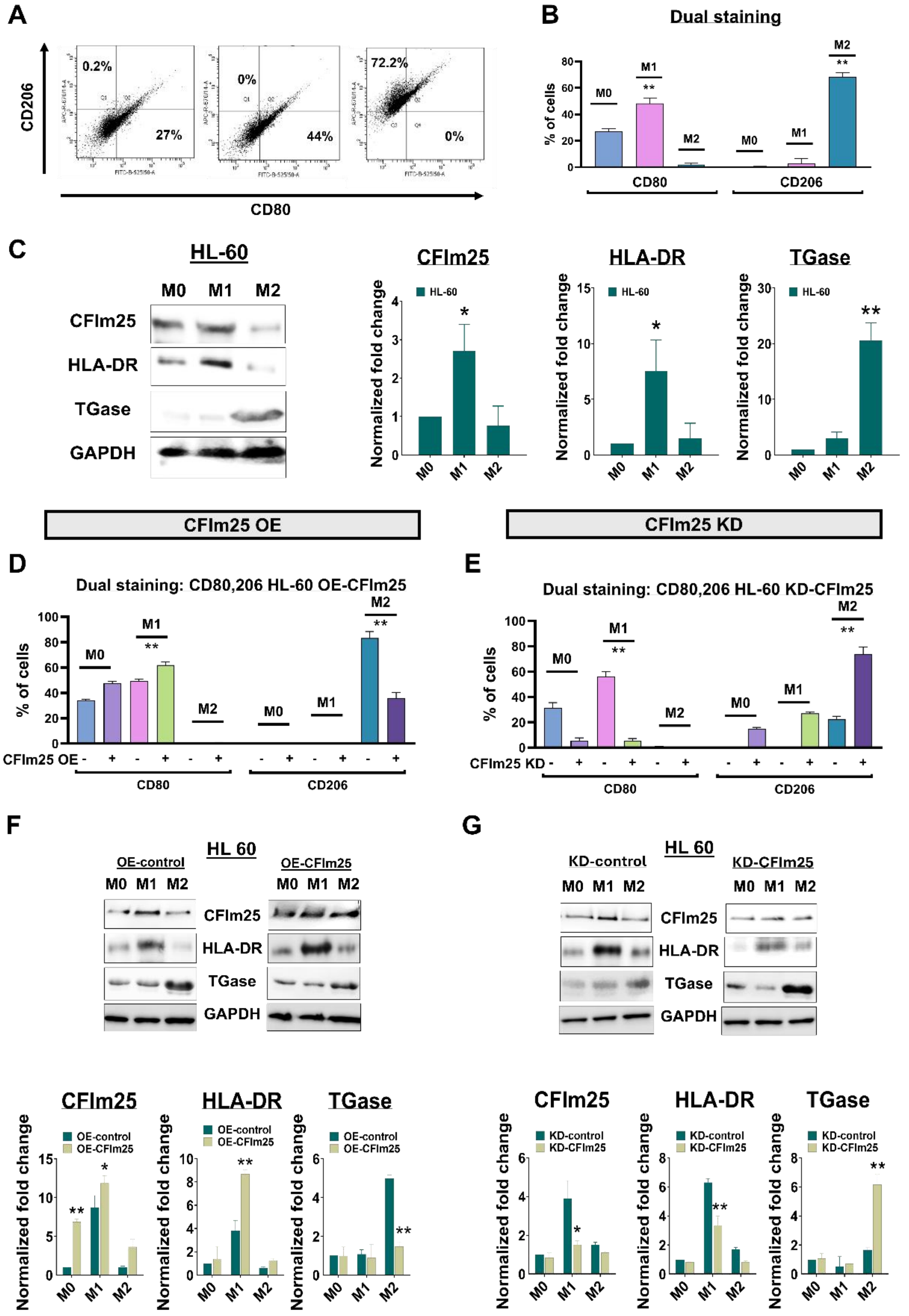


**Supplementary Figure 1. CFIm25 alteration drives macrophage surface marker expression in HL-60 cells (A)** Representative flow cytometry dot plots showing dual staining of M0, M1, and M2 polarized HL-60 cells for CD80 (M1 marker) and CD206 (M2 marker). Numbers indicate percentage of cells in each quadrant. **(B)** Quantification of flow cytometry data showing percentage of cells expressing CD80 and CD206 surface markers in M0, M1, and M2 polarized states. **(C)** Left: Western blot analysis of CFIm25, HLA-DR (M1 marker), and TGase (M2 marker) proteins in M0, M1, and M2 polarized HL-60 cells. GAPDH serves as loading control. Right: Densitometric quantification of protein levels normalized to M0 state. **(D)** Flow cytometry analysis of CD80 and CD206 expression in CFIm25-overexpressing (OE) HL-60 cells under M0, M1, and M2 polarizing conditions. **(E)** Flow cytometry analysis of CD80 and CD206 expression in CFIm25-knockdown (KD) HL-60 cells under M0, M1, and M2 polarizing conditions. **(F)** Top: Western blot analysis comparing control and CFIm25-overexpressing HL-60 cells for expression of CFIm25, HLA-DR, and TGase under different polarization conditions. GAPDH serves as loading control. Bottom: Densitometric quantification of protein levels normalized to PMA-treated respective controls (M0 state). **(G)** Top: Western blot analysis comparing control and CFIm25-knockdown HL-60 cells for expression of CFIm25, HLA-DR, and TGase under different polarization conditions. GAPDH serves as loading control. Bottom: Densitometric quantification of protein levels normalized to PMA-treated respective controls (M0 state). All data are shown as mean ± SEM from three independent experiments. $^{**}p < 0.01$, $^{*}p < 0.05$.

**Supplementary Figure 2. Macrophage polarization alters biochemical properties**

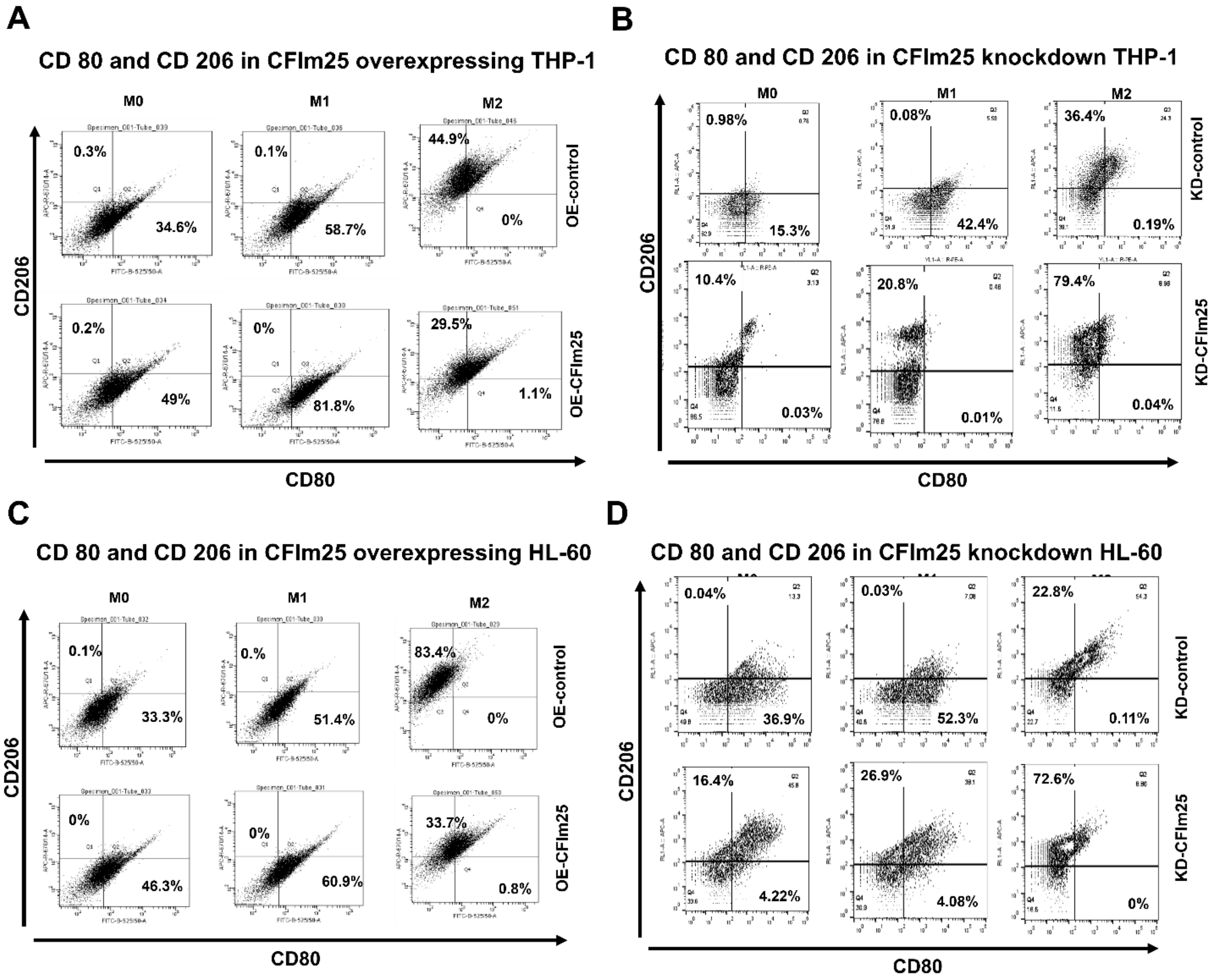


**Supplementary Figure 2. CFIm25 regulates macrophage surface marker expression. (A and B)** Representative flow cytometry dot plots showing dual staining for CD80 (M1 marker) and CD206 (M2 marker) in CFIm25-overexpressing (OE-CFIm25) THP-1 **(A)** or HL-60 **(B)** cells compared to control cells. Cells were analyzed in M0 (undifferentiated), M1, and M2 polarized states. **(C and D)** Representative flow cytometry dot plots showing dual staining for CD80 and CD206 in CFIm25-knockdown (KD-CFIm25) THP-1 **(C)** or HL-60 **(D)** cells compared to control cells. Analysis performed in M0, M1, and M2 polarized states. Numbers indicate percentage of cells in each quadrant.

**Supplementary Figure 3. Validation of efficient polarization by analysis of M1- or M2-specific biochemical activities and cytokine production**

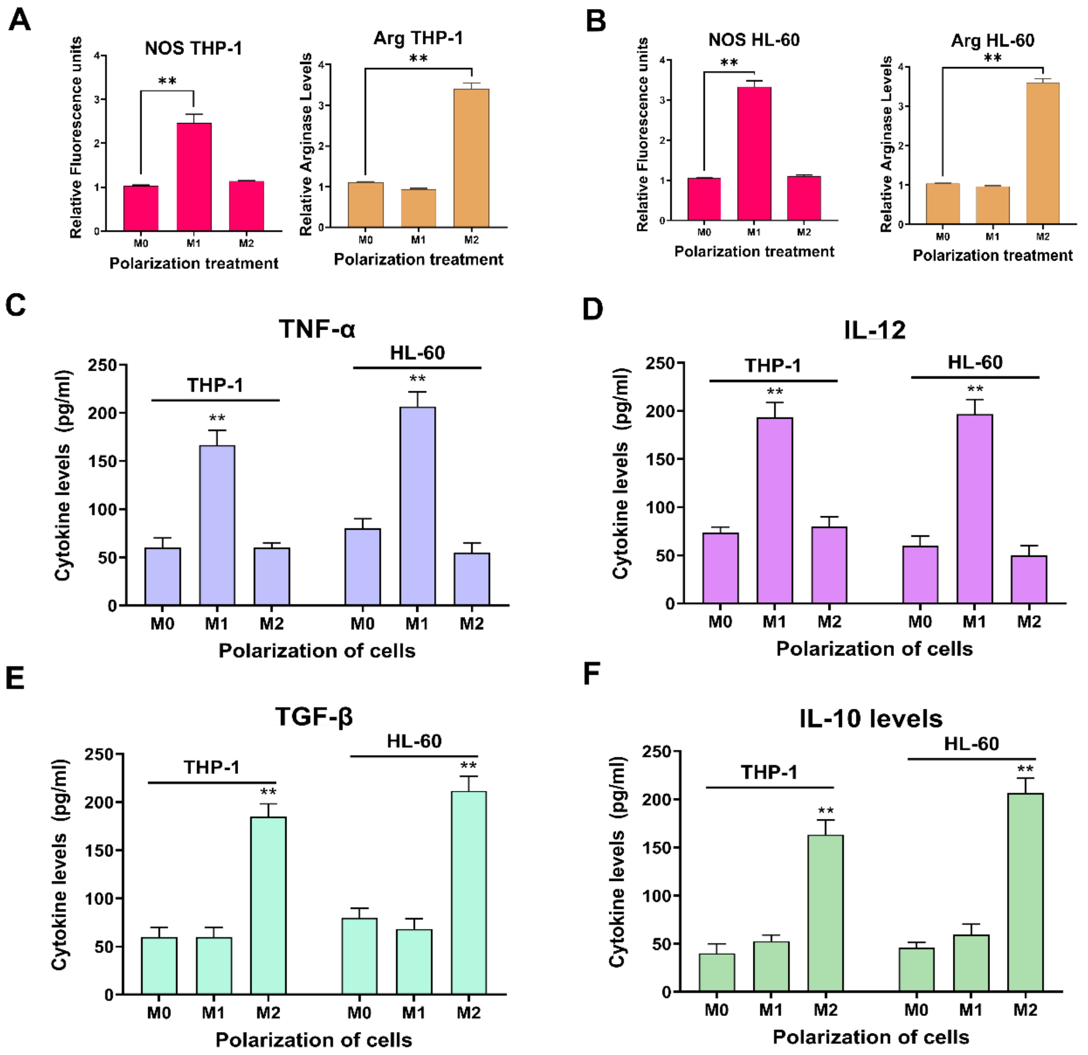


**Supplementary Figure 3. Validation of efficient polarization by analysis of M1- or M2-specific biochemical activities and cytokine production (A)** Analysis of THP-1 cells. Left: Nitric oxide synthase (NOS) activity measured by DAF-FM DA fluorescence in M0, M1, and M2 polarized cells. Right: Arginase activity assay in the same conditions. **(B)** Analysis of HL-60 cells. Left: NOS activity measured by DAF-FM DA fluorescence in M0, M1, and M2 polarized cells. Right: Arginase activity assay in the same conditions. For A-B, data are normalized to respective controls (M0 state) **(C)** TNF-α production measured by ELISA in M0, M1, and M2 polarized THP-1 and HL-60 cells. **(D)** IL-12 production measured by ELISA in M0, M1, and M2 polarized THP-1 and HL-60 cells. **(E)** TGF-β production measured by ELISA in M0, M1, and M2 polarized THP-1 and HL-60 cells. **(F)** IL-10 production measured by ELISA in M0, M1, and M2 polarized THP-1 and HL-60 cells. For C-F, data is represented as relative fluorescence units and are shown as mean ± SEM from three independent experiments $^{**}p < 0.01$, $^{*}p < 0.05$.

**Supplementary Figure 4. CFIm25 regulates macrophage biochemical properties in HL-60 cells**

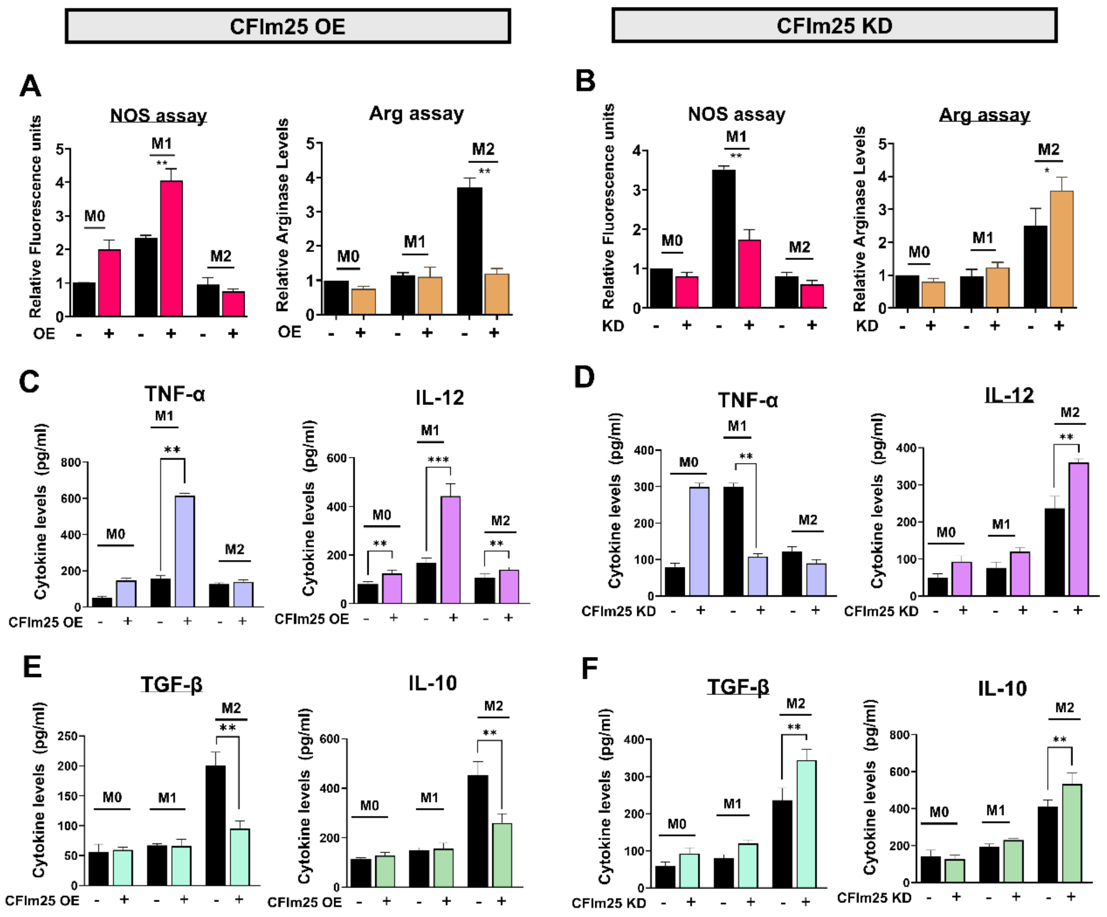


**Supplementary Figure 4. CFIm25 regulates macrophage biochemical properties in HL-60 cells. (A)** Analysis of Nitric oxide synthase (NOS) and Arginase (Arg) activity in CFIm25-overexpressing (OE) HL-60 cells. Left: NOS activity measured by DAF-FM DA fluorescence in M0, M1, and M2 polarized cells with or without CFIm25 OE. Right: Arginase activity assay in the same conditions. **(B)** Analysis of NOS and arginase in CFIm25-knockdown (KD) HL-60 cells. Left: NOS activity measured by DAF-FM DA fluorescence in M0, M1, and M2 polarized cells with or without CFIm25 KD. Right: Arginase activity assay in the same conditions. For A-B, data are normalized to respective controls (M0 state) **(C)** ELISA measurement of M1 cytokines in CFIm25 OE cells. Left: TNF-α levels in M0, M1, and M2 conditions. Right: IL-12 levels in the same condition. **(D)** ELISA measurement of M1 cytokines in CFIm25 KD cells. Left: TNF-α levels in M0, M1, and M2 conditions. Right: IL-12 levels in the same condition. **(E)** ELISA measurement of M2 cytokines in CFIm25 OE cells. Left: TGF-β levels in M0, M1, and M2 conditions. Right: IL-10 levels in the same conditions. **(F)** ELISA measurement of M2 cytokines in CFIm25 KD cells. Left: TGF-β levels in M0, M1, and M2 conditions. Right: IL-10 levels in the same conditions. For C-F, data is represented as relative fluorescence units and are shown as mean ± SEM from three independent experiments $^{**}p < 0.01$, $^{*}p < 0.05$.

**Supplementary Figure 5. Raw data for nitric oxide synthase (NOS) assay**

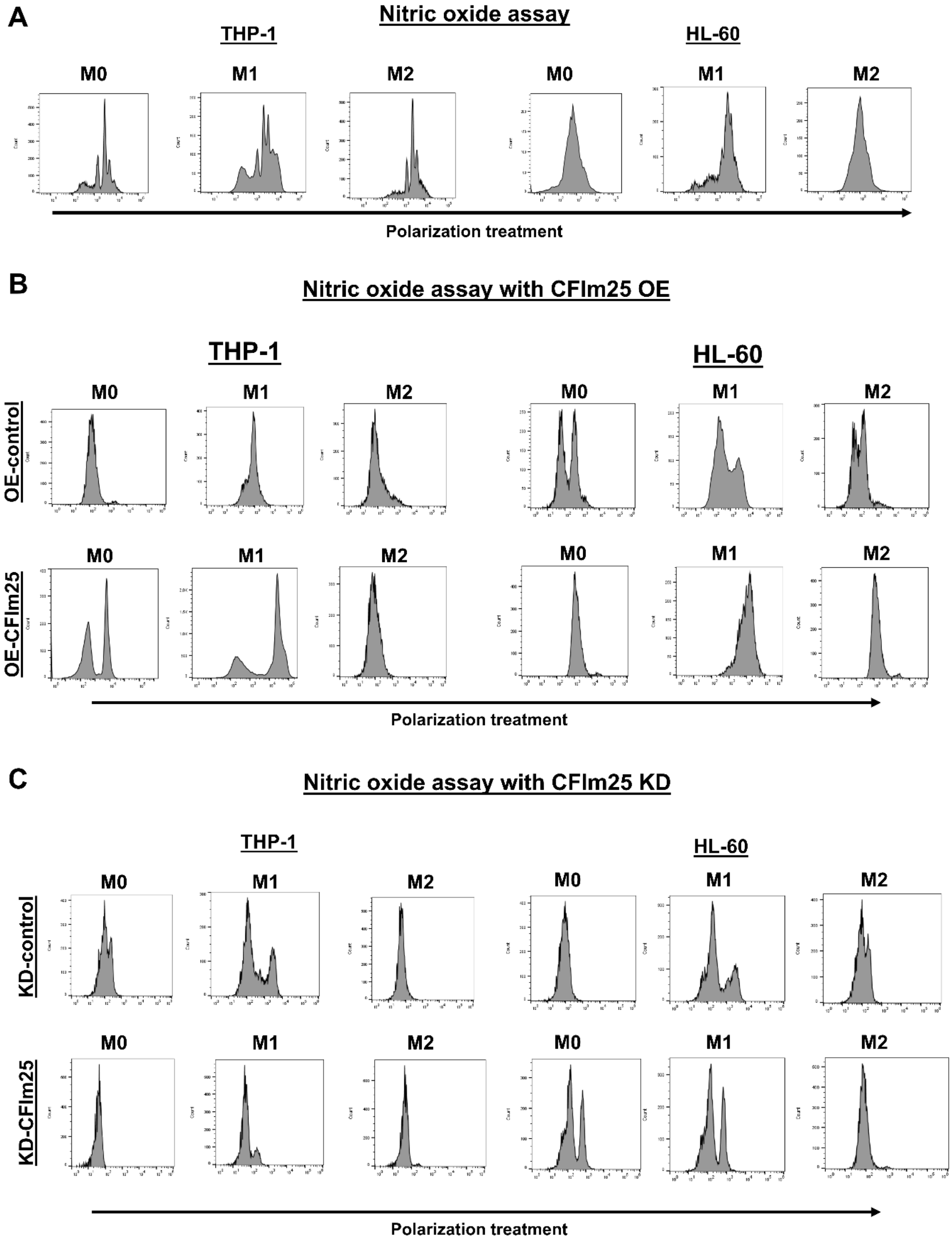


**Supplementary Figure 5. Raw data for nitric oxide synthase (NOS) assay**

**(A)** Representative flow cytometry histograms showing nitric oxide production measured by DAF-FM DA fluorescence in THP-1 and HL-60 cells under M0, M1, and M2 polarization conditions. **(B)** Representative flow cytometry histograms comparing nitric oxide production in CFIm25-overexpressing (OE-CFIm25) and control (OE-control) cells under different polarization conditions. Top panels show control cells and bottom panels show CFIm25 OE cells for both THP-1 and HL-60 cell lines. **(C)** Representative flow cytometry histograms comparing nitric oxide production in CFIm25-knockdown (KD-CFIm25) and control (KD-control) cells under different polarization conditions. Top panels show control cells and bottom panels show CFIm25 KD cells for both THP-1 and HL-60 cell lines.Data are representative of three independent experiments. Fluorescence intensity was measured by flow cytometry after staining with DAF-FM DA.

**Supplementary Figure 6. CFIm25 regulates core macrophage functions in HL-60 cells**

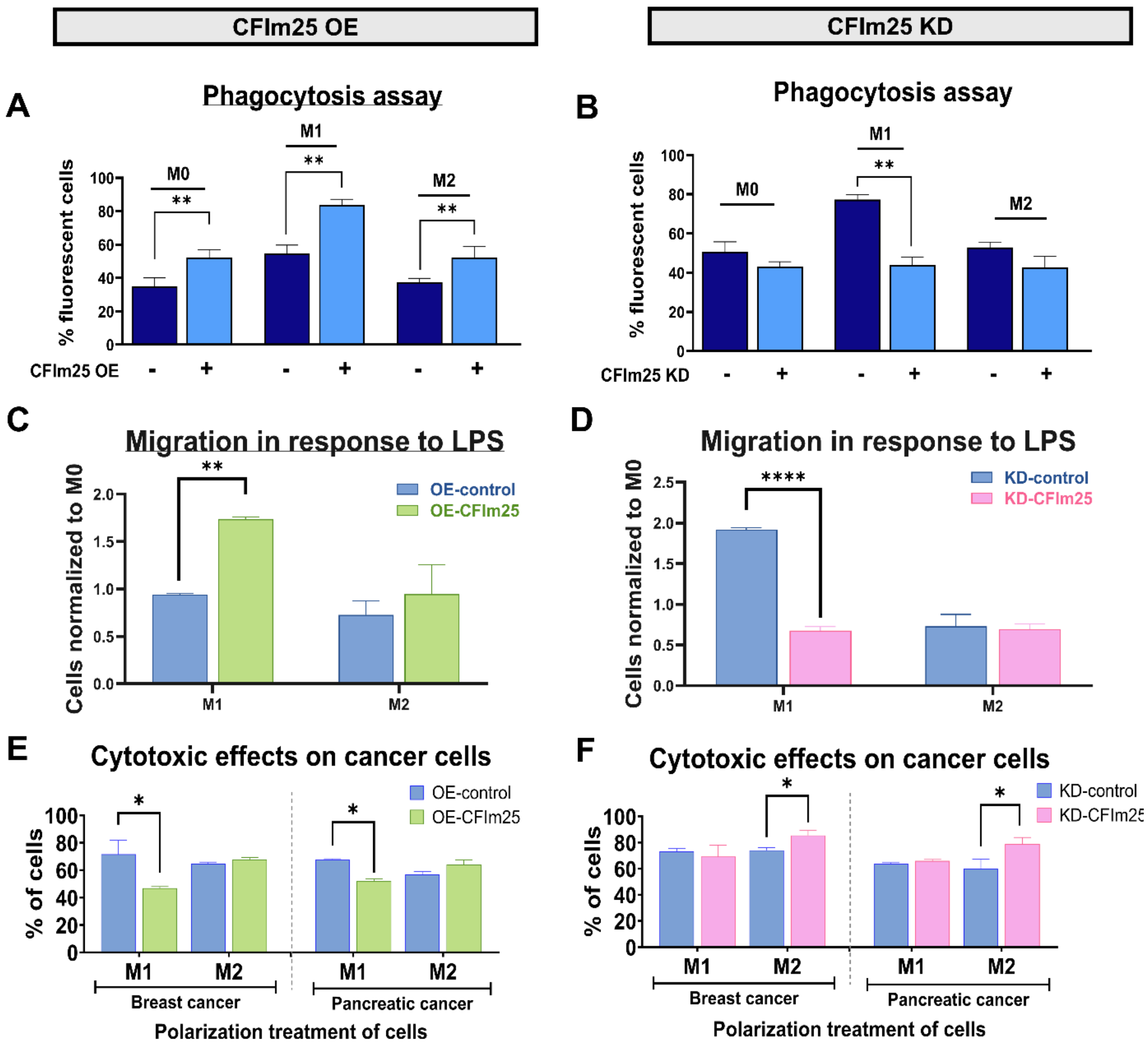


**Supplementary Figure 6. CFIm25 regulates core macrophage functions in HL-60 cells (A)** Phagocytic activity in CFIm25-overexpressing (OE) HL-60 cells. Flow cytometry analysis of fluorescently labeled IgG-coated latex bead uptake in M0, M1, and M2 polarized cells with or without CFIm25 OE. Data shown as percentage of fluorescent cells, mean ± SEM from three independent experiments. $^{**}p < 0.01$. **(B)** Phagocytic activity in M0, M1, and M2 polarized HL-60 cells with or without CFIm25 knockdown (KD). Data shown as percentage of fluorescent cells, mean ± SEM from three independent experiments. $^{**}p < 0.01$. **(C)** Chemotactic migration assay for CFIm25 OE cells. Quantification of cell migration through Transwell membranes in response to LPS chemoattractant. Data normalized to M0 control cells, mean ± SEM from three independent experiments. $^{*}p < 0.05$. **(D)** Chemotactic migration assay for

CFIm25 KD cells. Data normalized to M0 control cells, mean ± SEM from three independent experiments. $**p < 0.01$, $***p < 0.001$. **(E)** Cytotoxic activity of CFIm25-overexpressing macrophages against cancer cells. Indirect co-culture assay assessing the viability of breast cancer (MDA-MB-231) and pancreatic cancer (Mia-PaCa2) cells cultured in the lower chamber, with M1 or M2 polarized macrophages—with or without CFIm25 overexpression—seeded in the upper chamber separated by a permeable membrane. **(F)** Cytotoxic activity of CFIm25-overexpressing macrophages against cancer cells. Indirect co-culture assay assessing the viability of breast cancer (MDA-MB-231) and pancreatic cancer (Mia-PaCa2) cells cultured in the lower chamber, with M1 or M2 polarized macrophages—with or without CFIm25 knockdown—seeded in the upper chamber separated by a permeable membrane. Data are presented as percentage of viable cancer cells (mean ± SEM, $n = 3$ independent experiments). $**p < 0.01$, $*p < 0.05$.

**Supplementary Figure 7. Raw data for NOS assay and dual staining after AMO treatment.**

**A**

**CD 80 and CD 206 in M1-polarized THP-1 treated with AMO**

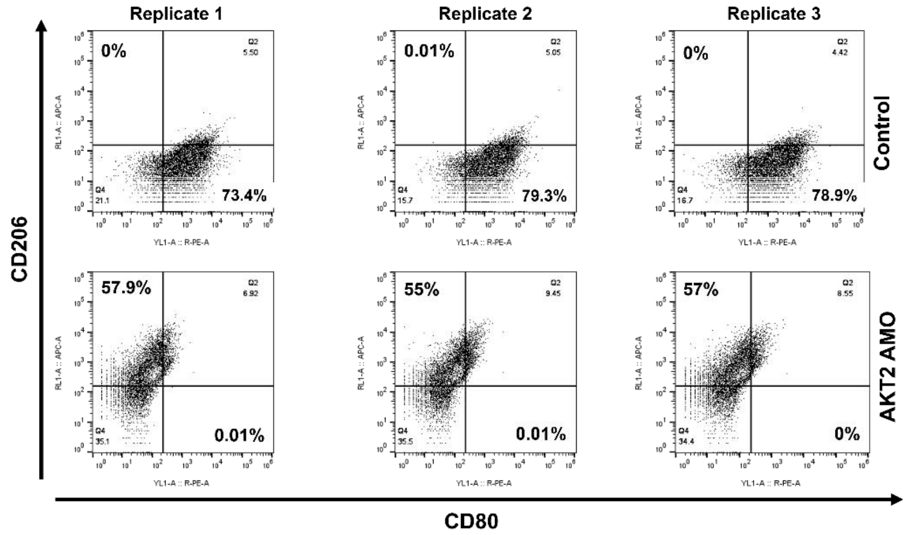


**B**

**Nitric oxide assay with M1 polarized THP-1 cells treated with AMO**

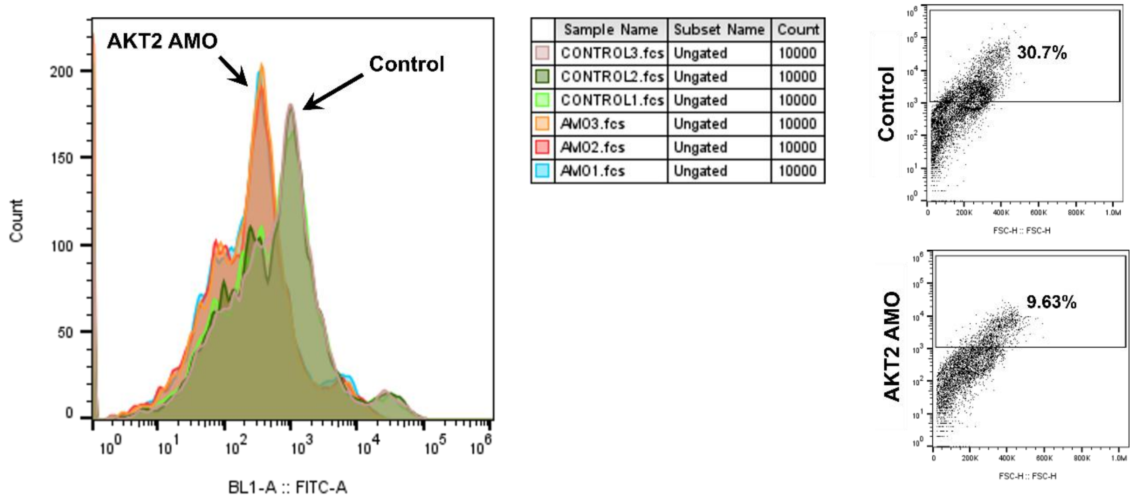


**Supplementary Figure 7. Raw data for NOS assay and dual staining after AMO treatment. (A)** *Left:* Merged flow cytometry histogram showing nitric oxide production measured by DAF-FM DA fluorescence in THP-1 M1 polarized cells treated with control or AKT2 AMO. *Right:* Representative flow cytometry dot blots comparing nitric oxide production in M1-polarized THP-1 cells with control or AKT2 AMO transfection. Numbers indicate percentage of cells in each quadrant. For all experiments, numbers indicate percentage of cells in each quadrant, and representative plots from three independent experiments are shown. **(B)** Flow cytometry dot plots showing dual staining for CD80 (M1 marker) and CD206 (M2 marker) in THP-1 M1 polarized cells treated with control or AKT2 AMO for three replicates.
